# Estimation of realized rates of genetic gain and indicators for breeding program assessment

**DOI:** 10.1101/409342

**Authors:** J.E. Rutkoski

## Abstract

Routine estimation of the rate of genetic gain (*ΔG*_*t*_) realized by a breeding program has been proposed as a means to monitor its effectiveness. Several methods of realized *ΔG*_*t*_ estimation have been utilized in other studies, but none have been objectively evaluated in a plant breeding context. Stochastic simulations of 80 rice (*Oryza sativa*) breeding programs over 28 years were done to generate data used to evaluate five methods of realized *ΔG*_*t*_ estimation in terms of error, precision, efficiency and correlation between true and predicted annual mean breeding values. Two indicators of *ΔG*_*t*_, the expected *ΔG*_*t*_ and the average number of equivalent complete generations (EqCg), were described and evaluated. At best, estimates of realized *ΔG*_*t*_ were over or underestimated by 15% and 27% when considering all 28 years and the past 15 years of breeding respectively. The best methods were the control population, estimated breeding value, and ERA trial methods. Among these, correlations between true and estimated *ΔG*_*t*_ were at best 0.59, indicating that these methods cannot very accurately rank breeding programs in terms of realized *ΔG*_*t*_. The expected *ΔG*_*t*_ and the average EqCg were shown to be useful indicators for determining if a non-zero genetic gain is expected. Determining which of the three best realized *ΔG*_*t*_ estimation methods evaluated, if any, would be appropriate for any given breeding program should be done with careful consideration of the objectives, resources, seed stocks, and structure of the data available.

## Introduction

Realized genetic gain is the change in average breeding value of a population over at least one cycle of selection for a particular trait or index of traits. Change in breeding or genetic values of populations over many cycles or years is referred to as genetic trend. When a genetic trend is linear, the rate of genetic gain per year, *ΔG*_*t*_, that is realized can be estimated by regressing the average breeding value on year (Eberhart, 1964). Assuming the breeding process remains unchanged and the trait of interest is quantitatively inherited according to the infinitesimal model (Fisher, 1918), this estimate is can be used to predict future genetic gain.

Analyses of genetic trends, often accompanied by estimation of realized *ΔG*_*t*_, have been conducted to analyze the outcomes of breeding programs or selection experiments in model species, livestock, and crops. In experimental populations of non-model and model species such as fruit fly (*Drosophila melanogaster*) and laboratory mouse (*Mus musculus*), selection experiments and assessments of genetic trends have been used to study the genetics of complex traits, selection limits, and to test quantitative genetic models and assumptions (Hill and Caballero, 1992). In livestock, reports of genetic trends are used to assess the effectiveness of breeding programs and to determine if adjustments are needed (Oldenbroek and van der Waaij, 2015). In crop species, estimation of realized *ΔG*_*t*_ is typically done after conducting selection experiments with the objective of comparing different breeding strategies (Hallauer et al., 2010) and to a lesser extent to assess the effectiveness of breeding programs. This may be because the adoption of varieties by growers in the targeted region has been considered to be the ultimate measure of success of the program (Brennan and Byerlee, 1991).

Various methods have been used to assess genetic trends depending on the species, data and germplasm that are available. Typically, for the analysis of genetic trends in model species, such as those described by Manning (1961), Kraaijeveld and Godfray (1997), and Swallow et al. (1998), an unselected control population is maintained and included in the phenotypic evaluations to be able to determine what changes in phenotypic value were due to artificial selection imposed by the experiment and what changes are due to environmental or other causes. In livestock breeding programs, because maintaining an unselected control population would be very costly, animal breeders assess genetic trends using estimated breeding values (EBVs), using methods reviewed by Garrick, 2010. EBVs are estimated using historical breeding program data and a mixed linear model with genetic covariance between relatives modeled according to additive relationships estimated using pedigrees. Then for each year, the average EBV for animals born in that year is computed and trends in average EBVs are analyzed. Genetic trends estimated in this way have been shown to be accurate under certain conditions (Sorensen and Kennedy, 1984) including accurate estimation of variance components and adequate connectedness across years. In crop selection experiments, analysis of genetic trends and estimation of realized *ΔG*_*t*_ is typically done by growing a sample of germplasm from each selection cycle in a common set of environments and regressing population means against year or cycle number (Eberhart, 1964). By growing all germplasm in one set of environments, trends in the average phenotypic values per-cycle can be attributed to genetic rather than environmental causes. In crop breeding programs on the other hand, samples of germplasm originating from each selection cycle are not usually available and so varieties released over time are used to represent the breeding germplasm. Such experiments are commonly referred to as ERA trials, popularized by Duvick who published a series of ERA trial studies between 1972 and 2004 to understand and quantify yield gains due to hybrid corn (*Zea mays*) breeding (reviewed by Duvick, 2005). In few studies, mixed models have been used to estimate realized *ΔG*_*t*_ in crop breeding programs using historical variety trial data (Mackay et al., 2011; Piepho et al., 2014).

Recently there has been significant interest in routinely analyzing genetic trends in order to estimate realized *ΔG*_*t*_ due to crop breeding programs to monitor their effectiveness (G.N. Atlin, personal communication, 2018). In order to do this effectively, careful planning and analysis is required to determine what data or germplasm should be archived or collected, and what experimental and statistical methods would be suitable given the resources and constraints of the breeding program. Potential methods of estimating realized *ΔG*_*t*_ should be evaluated based on their level of error and precision of realized *ΔG*_*t*_ estimation, and the resources they require. The ideal realized *ΔG*_*t*_ estimation method would be accurate, precise, and would utilize data already being generated by the breeding program.

Because estimation of realized *ΔG*_*t*_ requires that the performance of breeding germplasm be observed across time, in order for a breeding program to obtain an estimate of its realized *ΔG*_*t*_, the breeding program will ultimately need to have been operating for at least a few years or cycles of selection and it must have either kept samples of germplasm developed in different years or cycles, or accurate phenotypic data that is suitable for across-year analysis. In cases where these requirements are not met, indicators of realized *ΔG*_*t*_ may be useful for understanding if realized *ΔG*_*t*_ >0 is expected from the breeding effort. Indicators of realized *ΔG*_*t*_ could also be useful to monitor each time new parents are selected to help identify cases when the selection decisions will not lead to genetic gain. However, successful examples of this approach have not yet been reported.

In order to determine if a realized *ΔG*_*t*_ estimation method or indicator of realized *ΔG*_*t*_ is performing well, true values of realized *ΔG*_*t*_ should be known for comparison. Because true *ΔG*_*t*_ can only be known when data are from a simulated breeding program, computer simulation of plant breeding programs, reviewed by Sun et al., (2011) was used as a tool to evaluate realized *ΔG*_*t*_ estimation methods and indicators to determine which methods and indicators can be recommended for assessment of breeding programs and to understand their limitations. To identify the best method for estimation of medium to long term realized *ΔG*_*t*_, I evaluated a set methods similar to those previously reported for estimating medium to long term realized *ΔG*_*t*_ using 15 and 28 years of data generated from stochastic simulations of rice (*Oryza sativa*) breeding programs. Lastly, I examine potential indicators of realized *ΔG*_*t*_ that could be measured when obtaining reliable estimates of realized *ΔG*_*t*_ may not be immediately possible.

## Materials and methods

### Stochastic simulations for comparison of medium to long term ΔG_*t*_ estimation methods

A founder population of 140 individuals was simulated to have genome structure similar to that of elite *Indica* rice germplasm. To do so, 12 chromosomes of 110 centimorgans in length and an effective population size of 33 was assumed. The trait under selection was assumed to be quantitative, conferred by 1000 additive loci. To initiate the breeding program (Figure S1) founders underwent four generations of self-pollination. Thirty lines were selected as parents and intermated each year. Until at least two years worth of phenotypic data was available on new breeding lines, parents were randomly selected from the initial set of 140 inbred lines. After parents were selected, they were crossed in a half-diallel and then if at least two years worht of phenotypic data was available the top 100 F1s were selected, otherwise 100 F1s were selected at random. From the 100 F1s, 10 F2 progeny were generated per F1 and F2s were selfed for three more generations, resulting in 1000 F5s.

The 1000 F5s entered the testing and advancement process which followed one of two possible schemes (Table 1). In scheme A, 1000 F5s were phenotyped in location one with one replicate, and then the best 100 were selected for advancement. These 100 were phenotyped the following year at locations one through two with two replicates per location. From these 100, the best 50 were selected for advancement and phenotyped the following year at locations one through four with two replicates per location. In scheme B, 1000 F5s were phenotyped in location one with one replicate, and then the best 267 were selected for advancement. The 267 lines were phenotyped in the following year at locations two through four with one replicate per location. In both schemes, advancement from one stage to the next and selection of parents was restricted by family such that no more than three lines could be selected per family. In both schemes, once parent selection was based on performance and not at random, in the same year that parents were selected, parental lines were phenotypically evaluated with the same number of locations and replications as the previous stage of testing.

**Table 1:** Testing and advancement schemes followed in simulated breeding programs.

| Scheme | Stage | Lines tested | Locations | Replicates |
| --- | --- | --- | --- | --- |
| A | 1 | 1000 new | 1 | 1 |
|  | 2 | 100 from stage 1 | 1-2 | 2 |
|  | 3 | 50 from stage 2 | 1-4 | 2 |
|  | 4 | 30 current parent selections† | 1-4 | 2 |
| B | 1 | 1000 new | 1 | 1 |
|  | 2 | 267 from stage 1 | 2-4 | 1 |
|  | 3 | 30 current parent selections | 2-4 | 1 |

†Parent selections were made each year among all lines tested in the past three and two years for schemes A and B respectively

In both schemes, selections were based on best linear unbiased prediction BLUP assuming genotypes were independent and identically distributed (i.i.d.). For advancement, only phenotypic data from the current year was utilized whereas for parent selection, phenotypic data from the last three years and last two years were utilized in scheme A and B respectively. After the first round of the testing and advancement process, the best five lines were selected as checks that were henceforth phenotyped along with breeding lines at each stage of testing. Simulation of phenotypic data followed one of two different sets of assumptions of variance components (Table 2) in order to simulate high and low levels of heritability. The first stage of testing was always assumed to have a higher error variance compared to the subsequent stages. To simulate either a negative or positive non-genetic trend, 0.1*x* was added to or subtracted from the phenotypic values each year where *x* is the year of the breeding program. In total there were eight different breeding program scenarios consisting of all possible combinations of testing scheme, heritability level, and non-genetic trend. Each breeding scenario was repeated 10 times for a total of 80 simulations.

**Table 2:** Variance component assumptions for low and high heritability breeding program scenarios.

| Variances | Heritability |  |
| --- | --- | --- |
|  | Low | High |
| Genetic | 1 | 1 |
| Additive genotype-by-location | 0.75 | 0.225 |
| Additive genotype-by-year | 0.5 | 0.15 |
| Random genotype-by-location | 0.75 | 0.225 |
| Random genotype-by-year | 0.5 | 0.15 |
| Residual, stage 1 | 1 | 0.3 |
| Residual, stage 2 | 0.25 | 0.075 |

### Comparison of breeding scenarios based on true ΔG_t_

To compare true *ΔG*_*t*_ from each breeding scenario per time period, average true breeding value per line-year for each simulation and time period was used to fit the mixed model

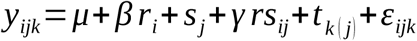

where *y*_*i*_ is the average true breeding value for line-year *i,* breeding scenario *j*, and simulation replicate *k, β* is a fixed regression coefficient for the average true *ΔG*_*t*_ across all simulations, *r*_*i*_ is a continuous covariate for the line-year *i, s*_*j*_ is a fixed factor for the breeding scenario *j, γ* is a fixed regression coefficient for the average true *ΔG*_*t*_ per breeding scenario, *rs*_*ij*_ is a fixed factor for the interaction between line-year *i* and breeding scenario *j, t* _*k* (*j*)_ is a random factor for the simulation replicate *k* nested within breeding scenario *j* with *t*_*k* (*j*)_ ∼ *N* (0, *σ*_*t*_ ^2^), and *ε*_*ijk*_ is the residual error with *ε*_*ijk*_ ∼ *N* (0, *Rσ*_*ε*_ ^2^). The diagonal of *R* contains a weighting factor to model a unique error variance for testing schemes A and B. Estimated marginal means and standard errors of linear trends were then then computed per breeding scenario. Statistical comparisons between pairs of trend estimates from different breeding scenarios were made using a Tukey’s multiplicity adjustment.

### Methods of medium to long term ΔG_t_ estimation

#### Control population

Using data exclusively from the final stage of testing, stage three in scheme A and stage two in scheme B, realized *ΔG*_*t*_ was estimated by fitting the mixed model:

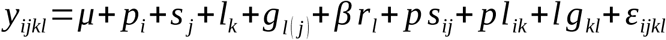

Where *y*_*ijlk*_ is the phenotypic value from population *i* observed in year *j* and location *k* on genotype *l, p*_*i*_ is the fixed effect of the *i*th population (either control, consisting of five long-term checks, or selected, consisting of the breeding lines), *s*_*j*_ is the random effect of the *j*th year with *s*_*j*_ ∼ *N* (0, *σ*_*s*_ ^2^), *l*_*k*_ is the fixed effect of the *k*th location, *g*_*l(j)*_ is the random effect of the *l*th genotype nested within the *j*th year with *g*_*l (j)*_ ∼ *N* (0, *σ*_*g*_ ^2^), *β* is a fixed regression coefficient for *ΔG*_*t*_, *r*_*l*_ is a continuous covariate for the line-year of genotype *l, p s*_*ij*_ is the random interaction effect between the *i*th population and the *j*th year with *ps*_*ij*_ ∼ *N* (0, *σ*_*ps*_ ^2^), *p l*_*ik*_ is the fixed interaction effect between the *i*th population and the *k*th location, *l g*_*kl*_ is the random interaction effect between the *k*th location and the *l*th genotype with *lg*_*kl*_∼ *N* (0, *σ*_*lg*_ ^2^), and *ε*_*ijkl*_ is the residual error with *ε*_*ijkl*_ ∼ *N* (0, *Rσ*_*ε*_ ^2^) where the diagonal of *R* contains the residual variances of *y*_*ijkl*_ to account for heterogeneous error variance.

#### EBV

After excluding terminal years of the breeding programs where phenotypic data from all testing stages was not present, breeding values were estimated based on the model

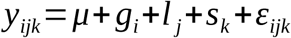

where *y*_*ijk*_ is the phenotypic value of genotype *i* observed in location *j* and year *k, g*_*i*_ is the effect of the *i*th genotype with *g*_*i*_ ∼ *N* (0, *A σ*_*g*_ ^2^) where *A* is the additive relationship matrix estimated using pedigree records from the simulated breeding program of interest, *l*_*j*_ is the effect of the *j*th location, *s*_*k*_ is the effect of the *k*th year, and *ε*_*ijk*_ is the residual error with *ε*_*ijk*_ ∼ *N* (0, *Rσ*_*ε*_ ^2^). The diagonal of *R* contains the residual variances of *y*_*ijk*_. Interactions between genotype and location and genotype and environment were not included due to software and hardware limitations. Next, the line-year, which is the year that a line entered the first stage of testing, was determined and the average EBV per line-year was computed. These average EBVs were then used as the response variable in the model:

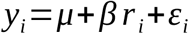

Where *y*_*i*_ is the average EBV for line-year *i, β* is a fixed regression coefficient for *ΔG*_*t*_, *r*_*i*_ is a continuous covariate for the line-year and *ε*_*i*_ is the residual error with *ε*_i_ ∼ *N* (0, *σ*_*ε*_ ^2^). The EBV method was repeated assuming different subsets of phenotypic data were available. When conducting statistical tests to compare methods, the EBV method applied to different phenotypic data subsets was considered as separate methods.

#### Variety trial

In each year of each simulated rice breeding program where phenotypic data on breeding lines was available prior to parent selection, the 30 lines selected as parents for crossing in the previous year were evaluated, making up stage four of Scheme A and stage three of Scheme B. A dataset similar to a variety trial dataset was then constructed by excluding data on lines that were never selected as parents, because only the top performing lines would be included in a variety trial. Then, according to Piepho et al. (2014), the simulated variety trial dataset was analyzed using the mixed model:

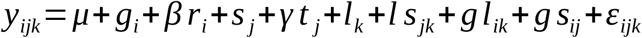

where *y*_*ijk*_ is the phenotypic value of genotype *i* observed in location *j* and year *k, g*_*i*_ is the random effect of the *i*th genotype with *g*_*i*_ ∼ *N* (0, *σ*_*g*_ ^2^), *β* is a fixed regression coefficient for *ΔG*_*t*_, *r*_*i*_ is a continuous covariate for the line-year of genotype *i, s*_*j*_ is the random effect of the *j*th year with *s*_*j*_ ∼ *N* (0, *σ*_*s*_ ^2^), *γ* is a fixed regression coefficient for the non-genetic gain or loss per year, *t* _*j*_ is a continuous covariate for the *j*th year, *l*_*k*_ is a random effect of the *k*th location with *l*_*k*_ ∼ *N* (0, *σ*_*l*_ ^2^), *l s*_*jk*_ is the random effect of the interaction between location and year with *ls* _*jk*_ ∼ *N* (0, *σ*_*ls*_ ^2^), *gl*_*ik*_ is a random effect of the interaction between genotype and location with *gl*_*ik*_ ∼ *N* (0, *σ*_*gl*_ ^2^), *g s*_*ij*_ is a random effect of the interaction between genotype and year with *g s*_*ij*_ ∼ *N* (0, *σ*_*gs*_ ^2^), and *ε*_*ijk*_ is the residual error with *ε*_*ijk*_ ∼ *N* (0, *Rσ*_*ε*_ ^2^), where the diagonal of *R* contains the residual variances of *y*_ijk_ to account for heterogeneous error variance.

#### ERA trial

For each line-year, genotypes belonging to that line-year were ranked according to their means calculated using phenotypic data from the final stage of testing, which was stage three of Scheme A and stage two of Scheme B. The top ranking genotypes per line-year were then selected for evaluation in a single simulated trial. The heritability level simulated was, 0.3, and one genotype was selected per line-year. Using the simulated trial data, the phenotypic mean for each line-year was then computed and then used as the response variable in the model:

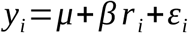

Where *y*_*i*_ is the phenotypic mean for line-year *i, β* is a fixed regression coefficient for *ΔG*_*t*_, *r*_*i*_ is a continuous covariate for the line-year and *ε*_*i*_ is the residual error with *ε*_*i*_ ∼ *N* (0, *σ*_*ε*_ ^2^).

#### Phenotype

Using phenotypic data from the final stage of testing, stage three of Scheme A and stage two of Scheme B, phenotypic means were computed per year and then used as the response variable in the model

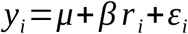

Where *y*_*i*_ is the phenotypic mean for year *i* within the final stage of testing, *β* is a fixed regression coefficient for *ΔG*_*t*_, *r*_*i*_ is a continuous covariate for the line-year and *ε*_*i*_ is the residual error with *ε*_*i*_ ∼ *N* (0, *σ*_*ε*_ ^2^).

### Comparison of medium to long term ΔG_t_ estimation methods

For two time periods, past 15 years and all 28 years, methods were compared based on their error, precision, efficiency and correlation between true annual mean breeding values and predicted annual mean breeding values. Error was defined as the absolute difference between the true and estimated realized *ΔG*_*t*_. True realized *ΔG*_*t*_ for each simulation and time period was estimated by fitting the mixed model

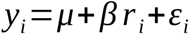

where *y*_*i*_ is the average true breeding value for line-year *i, β* is a fixed regression coefficient for true realized *ΔG*_*t*_, *r*_*i*_ is a continuous covariate for the line-year and *ε*_*i*_ is the residual error with *ε*_*i*_ ∼ *N* (0, *σ*_*ε*_ ^2^). The standard error of estimated realized *ΔG*_t_ was used as the measure of precision, efficiency was estimated as *p*_*t*_ +*n*_*t*_ / *N ×*100 where *p*_*t*_ is the number of true positives and *n*_*t*_ is the number of true negatives, and *N* is the total number of significance tests. True positives and true negatives were cases where the p-value of the true and estimated *ΔG*_*t*_ were both < 0.05 and both ≥ 0.05 respectively. For each of the metrics error, precision, efficiency, and correlation, means for each of the realized *ΔG_t_* estimation methods were compared by fitting the mixed model

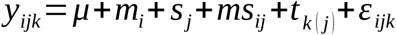

where *y*_*ijk*_ is the value of the metric of interest method *i,* breeding scenario *j*, and simulation replicate *k, m*_*i*_ is a fixed factor for realized *ΔG*_*t*_ estimation method *i, s*_*j*_ is a fixed factor for breeding scenario *j, ms*_*ij*_ is a fixed factor for the interaction between realized *ΔG*_*t*_ estimation method *i* and breeding scenario *j, t* _*k*(*j*)_ is a random factor for the simulation replicate *k* nested within breeding scenario *j* with *t*_*k*(*j*)_ ∼ *N* (0, *σ*_*t*_ ^2^), and *ε*_*ijk*_ is the residual error with *ε*_*ijk*_ ∼ *N* (0, *Rσ*_*ε*_ ^2^) the diagonal of *R* contains a weighting factor to model a unique error variance each *ΔG*_*t*_ estimation method *i*. For each metric, estimated marginal means per *ΔG*_*t*_ estimation method were compared using a Tukey’s multiplicity adjustment.

### Evaluation of indicators of ΔG_*t*_

#### Equivalent complete generations

The number of equivalent complete generations (EqCg), which is a measure of the equivalent number of discrete breeding cycles that have been completed in a breeding program where generations may overlap, was calculated per-line according to 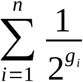 where *n* is the total number of known ancestors of the line, and *g*_*i*_ is the number of generations between the line and its ancestor *i* (Boichard et al., 1997). Only parents of crosses, as opposed to self-pollinations, were considered as ancestors. Furthermore, for any cross appearing in the pedigree, the individuals involved in that cross were counted as ancestors regardless of whether they had appeared in other crosses in the same pedigree. For example, a line that appears in a pedigree as both a parent and great-grandparent is counted as two different ancestors. Mean EqCg per line-year was then computed and trends in mean EqCg and the association of mean EqCg with average true breeding value was inspected and quantified within each simulated breeding program based on the Pearson’s correlation to determine if trend in EqCg could be used as an indicator of non-zero *ΔG*_*t*_.

#### Expected rate of genetic gain

For two time periods, past 15 years and all 28 years, the expected *ΔG*_*t*_ was computed each time parents were selected for crossing according to R/L where R is the expected response to selection in the current cycle which is the weighted average BLUP of the selected individuals minus the overall average BLUP and L is breeding cycle duration which is the weighted average age of the selected individuals. For the weighted averages, the relative contribution of a selected individual to the next generation was used as the weight. BLUPs were computed using all phenotypic data available within the time period of interest. Age of an individual was computed as the year of selection minus the year that the individual’s most recent parents were crossed. Self-pollinations were not considered as crosses. Average R/L was the overall estimate of the expected *ΔG*_*t*_. I considered average R/L as an additional method of realized *ΔG*_*t*_ estimation in some of the comparisons.

### Evaluation of the use of estimates of ΔG_*t*_ for comparing breeding programs

Associations between the expected *ΔG*_*t*_ or the estimated realized *ΔG*_*t*_ and true *ΔG*_*t*_ were inspected among the set of 80 simulated breeding programs to determine how well the methods of realized or expected *ΔG*_*t*_ could be used to compare breeding programs based on their true rates of *ΔG*_*t*_. The ability of the expected rate of genetic gain and estimated *ΔG*_*t*_ to accurately predict the true *ΔG*_*t*_ was quantified using the Pearson’s correlation, *r*. Absolute values of Pearson’s correlation greater than 0.22 were considered significant at an ? = 0.05 level of significance. To understand the cause of differences in levels of association between true and estimated *ΔG*_*t*_. Bias in estimated *ΔG*_*t*_ by scenario was inspected.

### Software

EBV estimation was implemented in REMLF90 (Misztal et al., 2014). All other analyses for the evaluation of medium to long term *ΔG*_*t*_ estimation methods were carried out using the R statistical programming language and environment (R Development Core Team, 2016) version 3.3.2. Stochastic simulations were carried out using the R package *BreedingSchemeLanguage* (Yabe et al., 2017). Mixed models analyses, other than EBV estimation, were implemented in the R package *lme4* (Bates et al., 2014). Tests for significant differences between means and linear trends were carried out using the packages *nlme* (Pinheiro et al., 2017) and *emmeans* (Lenth, 2018)

## Results

### Genetic trends and realized ΔG_t_ of simulated breeding programs

Genetic trends were linear for all eight scenarios (Figure 1). In all scenarios there was an initial period of seven years where there was no improvement in average breeding value because parents were selected at random from lines derived from the the initial population. Once selections could be made based on BLUP, annual improvements in average true breeding value were observed, however there was a year to year variability such that average true breeding value did not improve every year. This year to year variability was greater for scheme B compared to scheme A. Rates of realized *ΔG*_*t*_ in units of genetic standard deviations ranged from 0.19 to 0.32 (Table 3). The average rate of realized *ΔG*_*t*_ was 0.28 ± 0.0016 across all 28 years and 0.25 ± 0.0031 across the past 15 years. Within a scheme, high heritability led to higher rates of realized *ΔG*_*t*_ compared low heritability. When the non-genetic trend was negative and the heritability was low, scheme A lead to higher realized *ΔG*_*t*_ compared to scheme B in both time periods.

**Table 3:** True rates of realized *ΔG*_*t*_ for each breeding program scenario within two time periods.

| Scenario | True realized $\Delta G_t$ † | |
| --- | --- | --- |
|  | All 28 years | Past 15 years |
| Scheme A, + non-genetic trend, high heritability | 0.3 ± 0.0041 a | 0.28 ± 0.0059 a |
| Scheme A, + non-genetic trend, low heritability | 0.25 ± 0.0041 bc | 0.22 ± 0.0059 c |
| Scheme A, - non-genetic trend, high heritability | 0.31 ± 0.0041 a | 0.28 ± 0.0059 a |
| Scheme A, - non-genetic trend, low heritability | 0.26 ± 0.0041 b | 0.25 ± 0.0059 b |
| Scheme B, + non-genetic trend, high heritability | 0.32 ± 0.005 a | 0.28 ± 0.011 ab |
| Scheme B, + non-genetic trend, low heritability | 0.24 ± 0.005 cd | 0.21 ± 0.011 c |
| Scheme B, - non-genetic trend, high heritability | 0.31 ± 0.005 a | 0.27 ± 0.011 ab |
| Scheme B, - non-genetic trend, low heritability | 0.23 ± 0.005 d | 0.19 ± 0.011 c |
| Mean | 0.28 ± 0.0016 | 0.25 ± 0.0031 |

† Within a time period, values joined by the same letter are not significantly different

**Figure 1:**
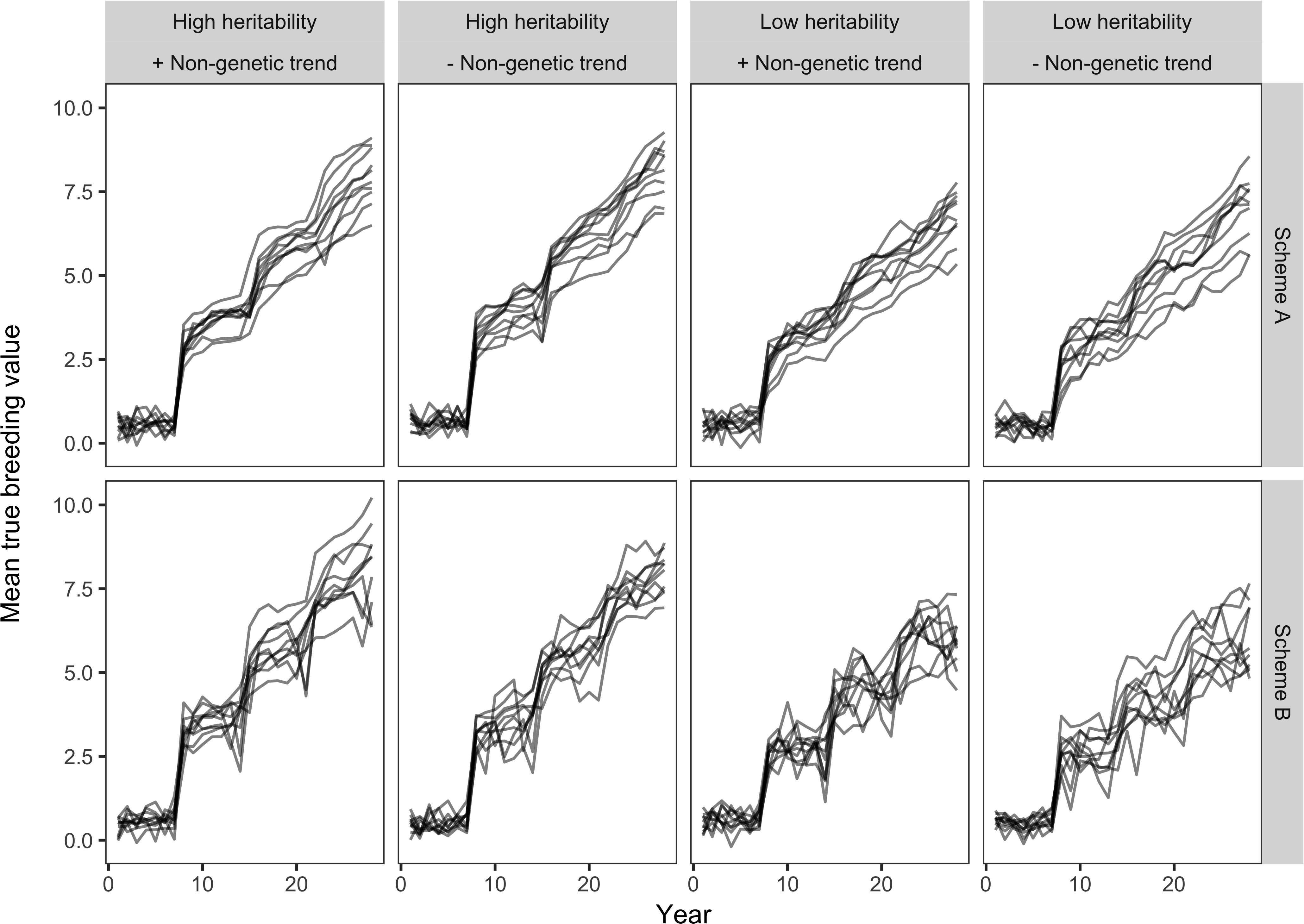
Trends in the mean true breeding value across years for each of the eight breeding scenarios simulated. Each panel corresponds to a different breeding scenario.

### Comparison of medium to long term realized ΔG_t_ estimation methods

Error of realized *ΔG*_*t*_ estimation methods (Table 4) ranged from 0.068 to 0.27 when only data from the past 15 years were utilized and from 0.043 to 0.23 when all 28 years were utilized. Considering that the average rate of gain was 0.25 in the past 15 year time period and 0.28 in the 28 year time period, these ranges in error correspond to an over or underestimation of realized *ΔG*_*t*_ by 27 to 108% in the 15 year time period and by 15 to 82% in the 28 year time period. Either the control population, EBV, and ERA trial methods had the lowest levels of error in the past 15 year time period. For the 28 year time period the control population method had the lowest level of error followed by the ERA trial, EBV, or variety trial method. Excluding checks from the dataset prior to using the EBV or variety trial methods led to a significant increase in error of these methods, with the error of the variety trial method affected to a greater extent. In fact, when the checks were excluded, the variety trial method had higher error than the phenotype method regardless of the time period. The best method that did not require that long-term checks be present in the breeding trial dataset was the ERA trial method, however, in the past 15 year time period the error of the ERA trial method was not significantly less than that of the phenotype method.

**Table 4:**
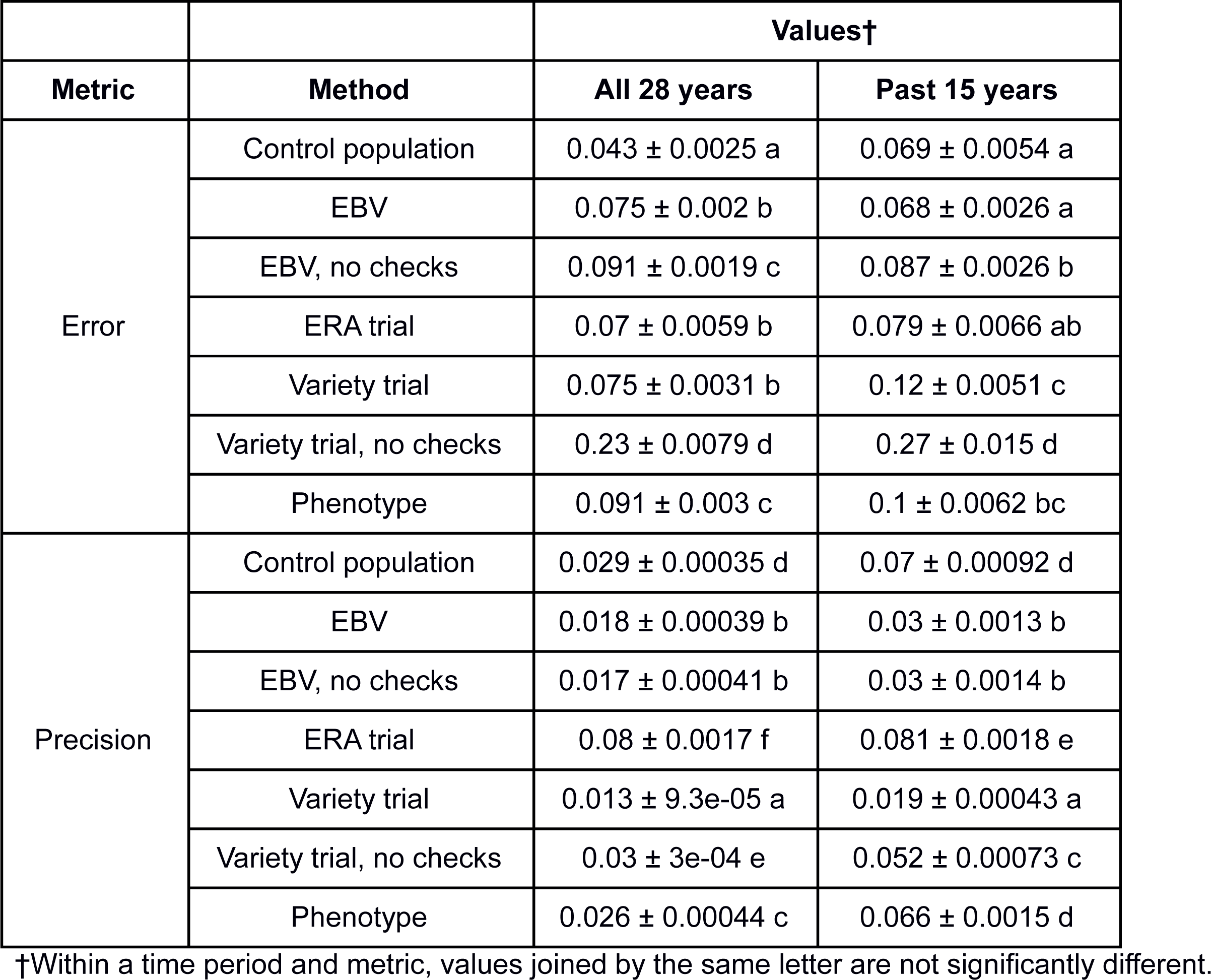
Average values of error and precision for each of the realized medium to long term *ΔG*_*t*_ estimation methods using data from two time periods.

|  |  | Values† |  |
| --- | --- | --- | --- |
| Metric | Method | All 28 years | Past 15 years |
| Error | Control population | 0.043 ± 0.0025 a | 0.069 ± 0.0054 a |
|  | EBV | 0.075 ± 0.002 b | 0.068 ± 0.0026 a |
|  | EBV, no checks | 0.091 ± 0.0019 c | 0.087 ± 0.0026 b |
|  | ERA trial | 0.07 ± 0.0059 b | 0.079 ± 0.0066 ab |
|  | Variety trial | 0.075 ± 0.0031 b | 0.12 ± 0.0051 c |
|  | Variety trial, no checks | 0.23 ± 0.0079 d | 0.27 ± 0.015 d |
|  | Phenotype | 0.091 ± 0.003 c | 0.1 ± 0.0062 bc |
| Precision | Control population | 0.029 ± 0.00035 d | 0.07 ± 0.00092 d |
|  | EBV | 0.018 ± 0.00039 b | 0.03 ± 0.0013 b |
|  | EBV, no checks | 0.017 ± 0.00041 b | 0.03 ± 0.0014 b |
|  | ERA trial | 0.08 ± 0.0017 f | 0.081 ± 0.0018 e |
|  | Variety trial | 0.013 ± 9.3e-05 a | 0.019 ± 0.00043 a |
|  | Variety trial, no checks | 0.03 ± 3e-04 e | 0.052 ± 0.00073 c |
|  | Phenotype | 0.026 ± 0.00044 c | 0.066 ± 0.0015 d |

†Within a time period and metric, values joined by the same letter are not significantly different.

Precision, measured using the standard error, of realized *ΔG*_*t*_ estimation methods (Table 4) ranged from 0.019 to 0.081 when using data from the past 15 years and from 0.013 to 0.08 when using data from all 28 years. The variety trial method was the most precise method followed by the EBV method. Excluding the checks from the dataset significantly reduced the precision obtained for the variety trial method. The least precise method in both time periods was the ERA trial method.

Efficiency of the *ΔG*_*t*_ estimation methods (Table S2) ranged from 79 to 100% when using data from the past 15 years and from 78 to 100% when using data from all 28 years. When using data from the past 15 years there was greater variability in efficiencies among the different *ΔG*_*t*_ estimation methods. Regardless of whether checks were included, the EBV and variety trial methods had efficiency levels of at least 98%. The next most efficient method in this time period was the control population method followed by the phenotype and ERA trial methods. When using data from all 28 years, all methods had 100% efficiency except for the ERA trial method which had an efficiency level of 78%.

Correlations between the annual true mean breeding values and the predicted annual mean breeding values (Table S2) were at least 0.9 with the exception of three cases. When using data from the past 15 years, correlations were 0.74 and 0.51 for the phenotype and ERA trial methods respectively and when using data from all 28 years, the ERA trial method had a correlation of 0.49.

### Impact of incomplete data on the effectiveness of the EBV method

Accuracies of the EBV method excluding different subsets of data ranged from 0.052 to 0.11 in the last 15 year time period and 0.057 to 0.1 in the 28 year time period (Table 5). Excluding data on lines discarded in stage one either improved or had no impact on error. Excluding data on the last testing stage or the checks led to a significant reduction in error, except when data was also excluded on the lines discarded in stage one.

**Table 5:**
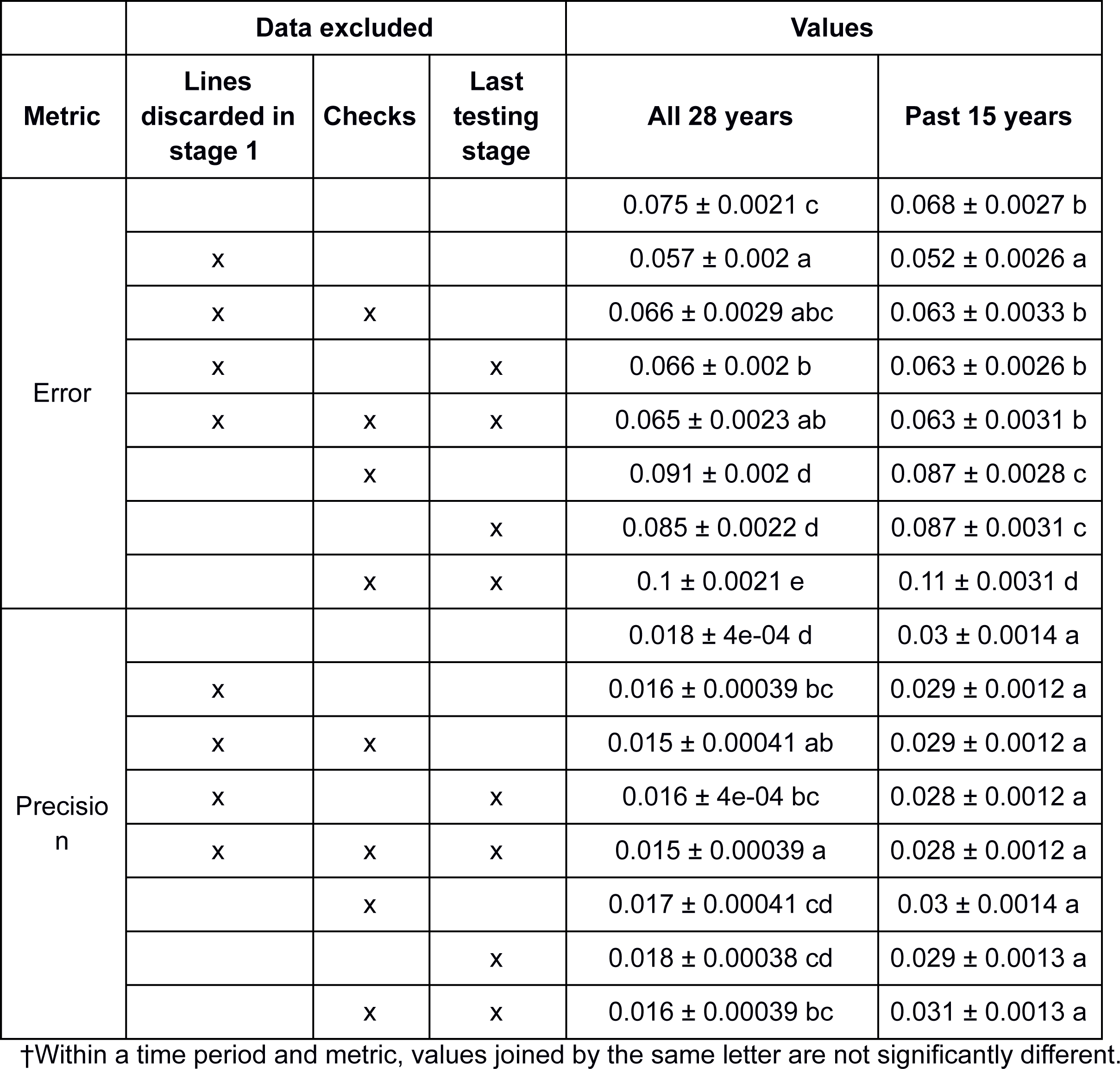
Average values of error and precision for the EBV method of *ΔG*_*t*_ estimation using data from two time periods and excluding different sets of data.

|  | Data excluded |  |  | Values |  |
| --- | --- | --- | --- | --- | --- |
| Metric | Lines discarded in stage 1 | Checks | Last testing stage | All 28 years | Past 15 years |
| Error |  |  |  | 0.075 ± 0.0021 c | 0.068 ± 0.0027 b |
|  | x |  |  | 0.057 ± 0.002 a | 0.052 ± 0.0026 a |
|  | x | x |  | 0.066 ± 0.0029 abc | 0.063 ± 0.0033 b |
|  | x |  | x | 0.066 ± 0.002 b | 0.063 ± 0.0026 b |
|  | x | x | x | 0.065 ± 0.0023 ab | 0.063 ± 0.0031 b |
|  |  | x |  | 0.091 ± 0.002 d | 0.087 ± 0.0028 c |
|  |  |  | x | 0.085 ± 0.0022 d | 0.087 ± 0.0031 c |
|  |  | x | x | 0.1 ± 0.0021 e | 0.11 ± 0.0031 d |
| Precision |  |  |  | 0.018 ± 4e-04 d | 0.03 ± 0.0014 a |
|  | x |  |  | 0.016 ± 0.00039 bc | 0.029 ± 0.0012 a |
|  | x | x |  | 0.015 ± 0.00041 ab | 0.029 ± 0.0012 a |
|  | x |  | x | 0.016 ± 4e-04 bc | 0.028 ± 0.0012 a |
|  | x | x | x | 0.015 ± 0.00039 a | 0.028 ± 0.0012 a |
|  |  | x |  | 0.017 ± 0.00041 cd | 0.03 ± 0.0014 a |
|  |  |  | x | 0.018 ± 0.00038 cd | 0.029 ± 0.0013 a |
|  |  | x | x | 0.016 ± 0.00039 bc | 0.031 ± 0.0013 a |

†Within a time period and metric, values joined by the same letter are not significantly different.

Levels of precision of the EBV method when using different subsets of data (Table 5) ranged from 0.028 to 0.031 in the past 15 year time period and from 0.015 to 0.018 in the 28 year time period. None of the differences in precision were significant in the 15 year time period. In the 28 year time period the highest levels of precision were observed when data on the lines discarded in the stage one and the checks were excluded.

Efficiency of the EBV method was unaffected by the exclusion of different subsets of data (Table S3). In all cases 100% efficiency was observed. Similarly, correlations between the annual true mean breeding values and the predicted annual mean breeding values based on the EBV method were also mostly unaffected by the subset of data used (Table S3). Correlations ranged from 0.97 to 0.98 in the past 15 year time period and from 0.97 to 0.99 in the 28 year time period.

### Evaluation of ΔG_t_ indicators

Trends in average EqCg (Figure 2) reflected the trends in average true breeding values (Figure 1). Within a breeding program, the average correlation between mean EqCg per line-year and mean true breeding value per line-year was 0.98 ± 0.0010 over all 28 years and 0.98 ± 0.001 over the last 15 years. Average R/L had an error level of 0.079 ± 0.0033 in the past 15 year time period which was not significantly different from the control population, EBV, and ERA trial methods. In the 28 year time period average R/L had an error of 0.064 ± 0.0029 which was less accurate than the control population method but just as accurate as the EBV, ERA trial, and variety trial methods.

**Figure 2:**
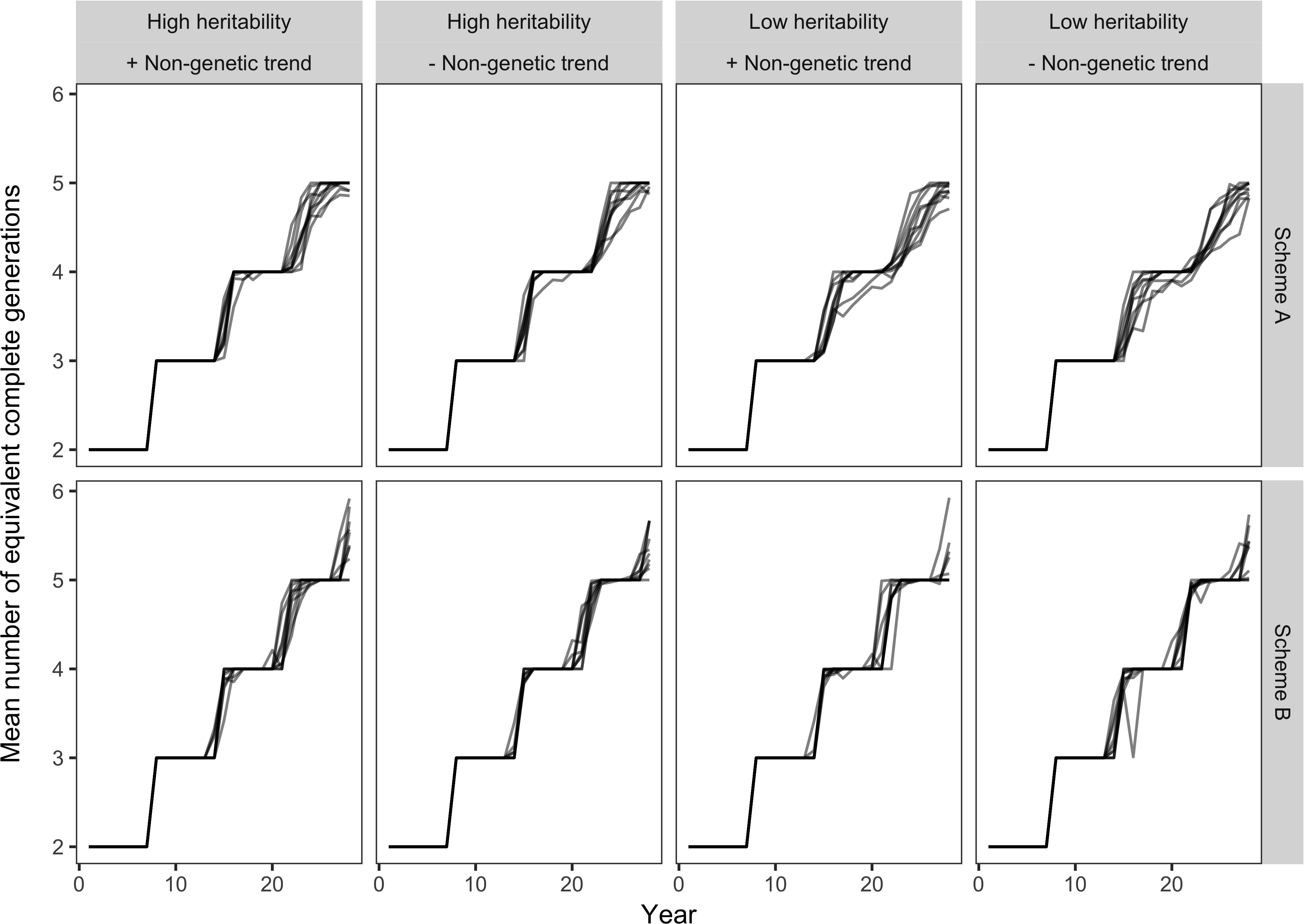
Trends in the average number of equivalent complete generations across years for each of the eight breeding scenarios simulated. Each panel corresponds to a different breeding scenario.

### Association between the true realized ΔG_t_ and estimated or expected realized ΔG _t_

Positive linear relationships between the true realized *ΔG*_*t*_ and estimated or expected realized *ΔG*_*t*_ were observed for the average R/L, control population, EBV, ERA trial, and phenotype methods in the 15 year (Figure 3) time period. This observation was consistent in the 28 year time period (Figure S2), except that estimated realized *ΔG*_*t*_ based on the EBV method was not associated with true realized *ΔG*_*t*_. Interestingly, in the 28 year time period the variety trial method with data on checks excluded provided estimates of realized *ΔG*_*t*_ which were significantly negatively correlated with the true realized *ΔG*_*t*_. The best method based on association between true realized *ΔG*_*t*_ and estimated or expected realized *ΔG*_*t*_ was the control population method in the 15 year time period and the expected gain method based on average R/L in the 28 year time period. In the 28 year time period the control population method was second best. Systematic error (bias) in expected or estimated realized *ΔG*_*t*_ was detected. For example among the methods that lead to the highest associations between true and expected or estimated realized *ΔG*_*t*_ in the 15 year time period, bias was associated with the breeding scenario (Figure 4). This association was most pronounced for the EBV method and least pronounced for the ERA trial method.

**Figure 3:**
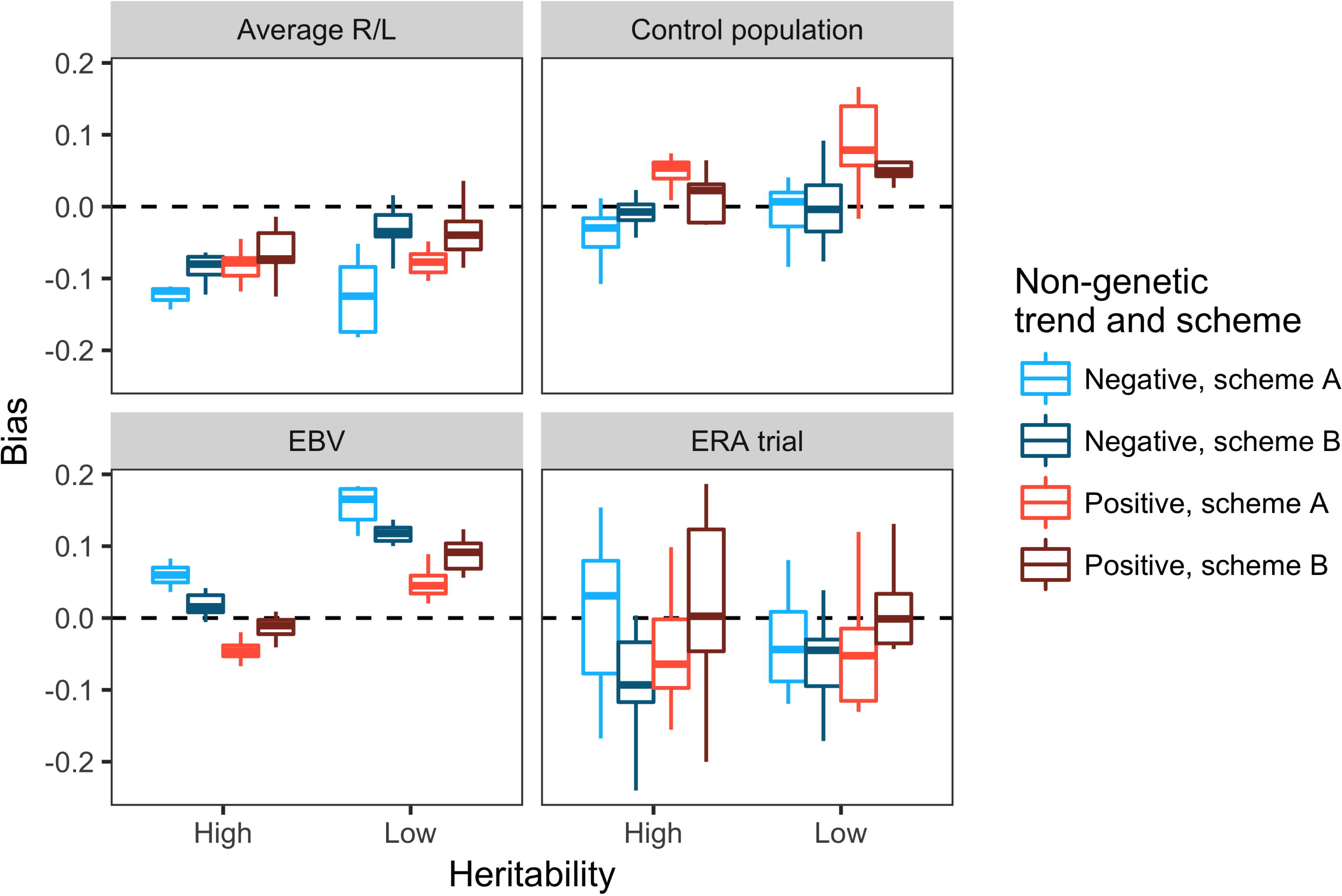
Scatterplots showing the relationships between estimated and true *ΔG*_*t*_ in the past 15 year time period for different methods of *ΔG*_*t*_ estimation. Each point within a panel represents one simulated breeding program. Pearson’s correlations, *r*, between estimated and true *ΔG*_*t*_ are shown in the lower right corner of each panel.

**Figure 4:**
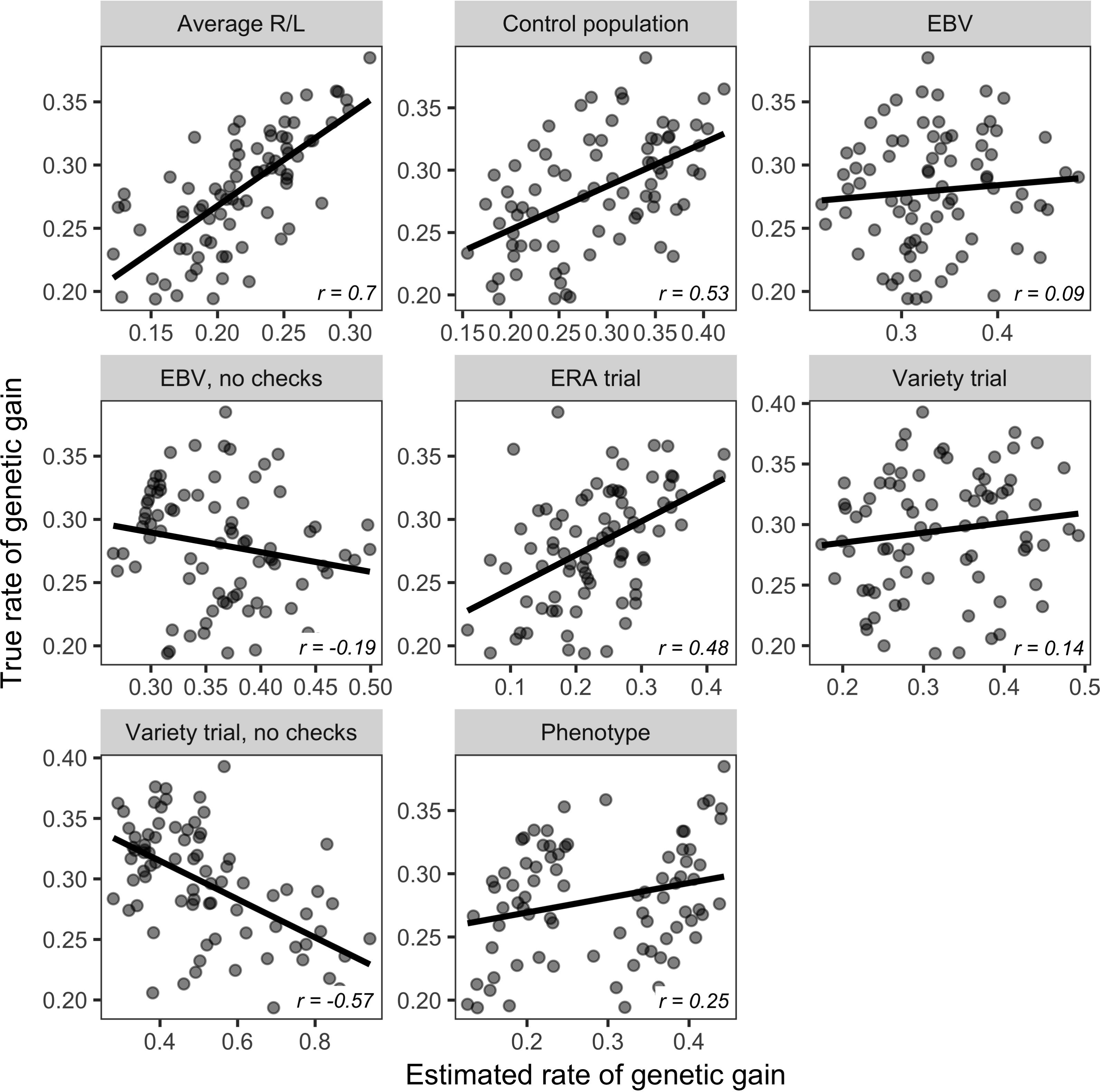
Boxplots showing the bias of the estimates of *ΔG*_*t*_ for each breeding scenario and for four methods of *ΔG*_*t*_ estimation in the past 15 year time period. Dotted lines are drawn at zero. Blue and red boxes represent scenarios where the non-genetic trend was negative and positive respectively. Light blue and red boxes correspond to scenarios where testing scheme A was used. Dark red and blue boxes correspond to scenarios where testing scheme B was used. Low heritability scenarios are on the right of each panel, whereas high heritability scenarios are on the left.

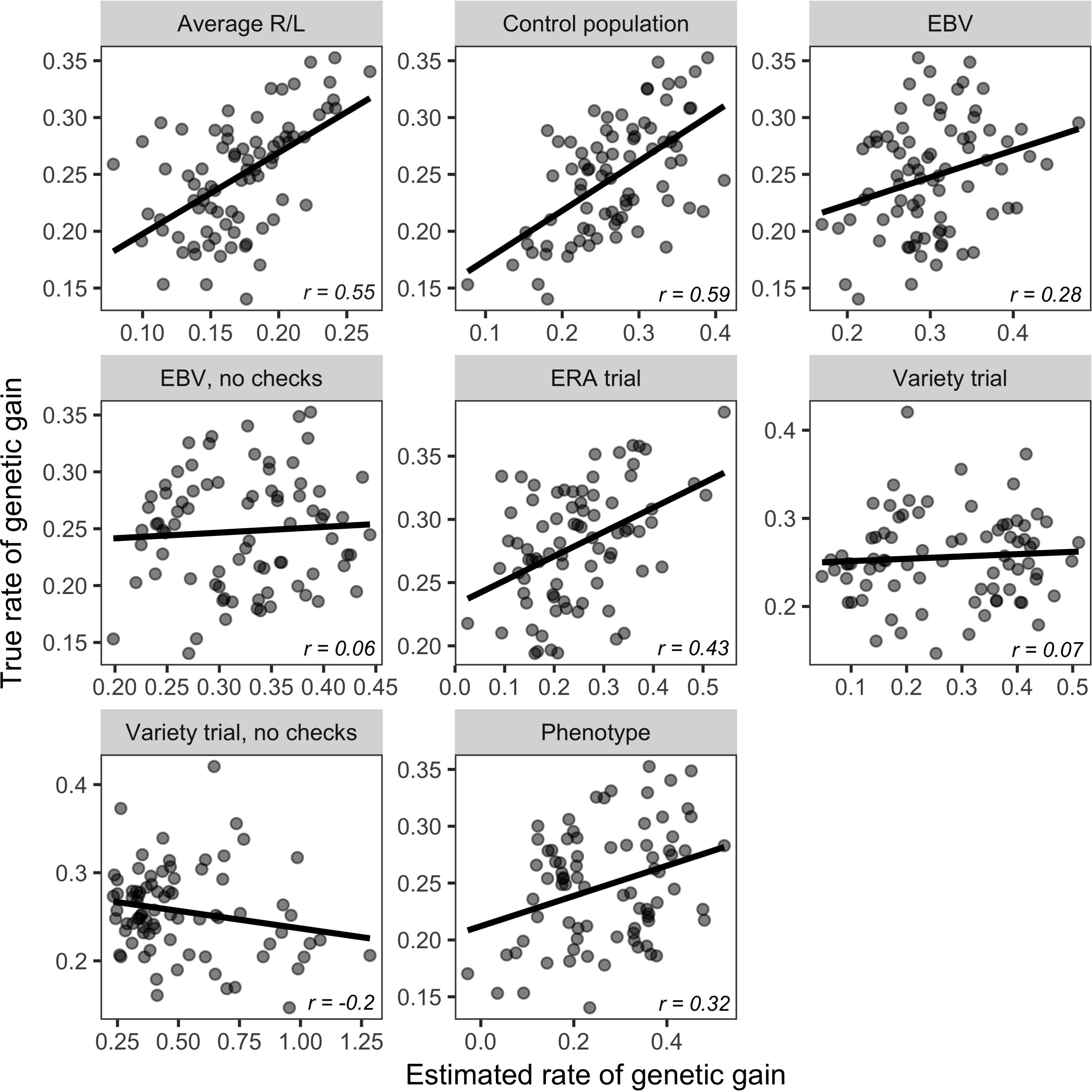

## Discussion

### Simulated breeding scenarios

The breeding scenarios simulated in this study were designed to mimic actual medium to low budget breeding programs of rice and other self-pollinated crops. As in real breeding programs, the testing and selection practices imposed in the simulations may not have been ideal for maximizing *ΔG*_*t*_, however all simulated breeding scenarios lead to a linear increase in true breeding values over time. Testing scheme A lead to less variability in genetic trend compared to testing scheme B. This is probably because with testing scheme B there was more disconnectivity in the data leading to confounding of breeding value, year, and location effects, reducing my ability to achieve the expected level of selection error on a consistent basis. The stair-step pattern in the genetic trend observed (Figure 1) especially in the earlier years of the breeding programs was not surprising. This occurred because at the outset of the breeding program generations were mostly discrete and waves of improved material appear corresponding to the completion of a complete breeding cycle. The stair-step pattern diminishes towards the end of the simulations because the generations begin to overlap to a greater extent. This phenomena can also be observed upon inspection of the trends EqCg. Change in EqCg becomes more continuous towards the end of the breeding programs. In real breeding programs, it should be kept in mind that when a breeding program is first initiated, there may be a period of time where genetic improvement will come in waves and not a follow a smooth linear trend. In other words, a measurable improvement in the population is not to be expected every year, especially if a breeding program was recently initiated.

### Estimation of medium to long term realized ΔG_t_

Error of realized *ΔG*_*t*_ estimation was considered to be the most important factor when comparing the effectiveness of different methods because error indicates how close the estimates are from the true values of realized *ΔG*_*t*_. Error of different methods varied to a large extent, thus it was useful for discriminating among methods. Precision, based on the standard error of realized *ΔG*_*t*_ estimates, was considered an important metric, however evaluating methods based on precision alone would be misleading because methods with high levels of precision often had unacceptable levels of error. Often in assessment of statistical methods in real datasets precision is used as a primary indicator of a method’s performance, but based on what was observed, evaluating realized *ΔG*_*t*_ estimation methods based on precision alone is not sufficient. A method may provide an estimate of realized *ΔG*_*t*_ which is highly precise but highly inaccurate. Efficiency was found to be a useful metric for discriminating between different realized *ΔG*_*t*_ estimation methods and it is considered important because significance thresholds are commonly used when determining if realized *ΔG*_*t*_ *>* 0. Lower levels of efficiency indicate that a method is more likely to misdiagnose the presence or absence of non-zero realized *ΔG*_*t*_ based on the p-value of the estimate. As with precision, methods with high levels of efficiency sometimes had low levels of error, and so evaluation of methods based on efficiency alone is not recommended. Finally, correlation with the average true breeding values over time is important if one is interested in predicting and visualizing the population means over time. It should be noted that the control population and variety trial methods only predicted the population mean breeding value based on a the linear rate of realized *ΔG*_*t*_ estimated, so these method do not provide a visual showing how the population means over time deviate from linearity. Conveniently, correlation was associated with efficiency, so it was not difficult to identify efficient methods that also work well for predicting the actual population means per year.

Considering all the metrics evaluated, the best method for estimation of realized *ΔG*_*t*_ in the 15 year time period was the EBV method either including data on long-term checks or excluding data on lines discarded after the first stage of testing. This method had the lowest level of error, the second highest level of precision, 100% efficiency, and the highest correlation between and true and estimated annual average breeding values (0.98). The main limitation to utilizing the EBV method in practice is that it requires pedigree records have been kept since the beginning of the breeding program so that additive relationships between lines can be estimated. The second factor potentially limiting the application of this method is the data structure required. There must be good connectivity between years for the EBV method to work well. Ideally the connectivity should be as good as or better than that of the data simulated in this study. Our results suggest that the use of long-term checks and discarding data on lines tested for only one year can help to increase the connectivity and the performance of the EBV method. Because it can be difficult to judge whether or not a dataset is suitable for estimation of realized *ΔG*_*t*_ based on the EBV method. The best way to do so may be to simulate the breeding and testing procedure used in the breeding program of interest assuming a realistic non-genetic trend, and then evaluate the effectiveness of the EBV method using the simulation data. The added benefit of this step would be that the expected rate of *ΔG*_*t*_ could also be estimated for comparison with that which was realized. Agreement between expected and realized *ΔG*_*t*_ would provide stronger evidence that the estimate of realized *ΔG*_*t*_ is accurate.

In the 28 year time period, the control population method was the best method for estimation of realized *ΔG*_*t*_ considering all the metrics evaluated. This method had the lowest error, moderately high precision, 100% efficiency, and high correlation between true and predicted annual average breeding values (0.97). The main limitation to utilizing this method in practice is that in order to estimate realized *ΔG*_*t*_ over long time periods it requires that the same check(s) have been phenotyped every year, a condition that is almost never met in data from real breeding programs. Furthermore, in real field experiments, checks that are grown for many years become increasingly susceptible to disease which can prevent meaningful phenotyping of other important traits and increase the magnitude of the non-genetic trend. While pathogen evolution can be a legitimate source of a non-genetic trend, if the objective is to measure realized *ΔG*_*t*_ for the trait of interest in the absence of disease, then loss of disease resistance in the checks would lead to an overestimation of realized *ΔG*_*t*_, leading to even higher levels of error than what was observed in this study. Due to these issues, the control population method for the estimation of realized *ΔG*_*t*_ over long time periods may not always be suitable, even if the data structure permits its use. The next best method in this time period was the EBV method excluding data on the lines discarded in the first stage of testing. Performance of this method was similar to that of the control population method except that it was slightly less accurate, but more precise.

Over shorter time periods, the control population method would be a good alternative to the EBV method assuming the same check(s) have been evaluated each year in the time period of interest. In the past 15 year time period of our experiment, the control population method was just as accurate as the EBV method. Precision was moderate to low relative to the other methods, but efficiency and correlation between true and estimated annual mean breeding values was 91% and 0.93 respectively which I consider to be acceptable. The advantage of using this method rather than the EBV method is that it is more flexible in that it can be used with datasets that are disconnected across years and it does not require that pedigree records have been kept from the beginning of the breeding program. As mentioned earlier, disadvantages of this method are that it cannot be used to estimate realized *ΔG*_*t*_ for the trait of interest in the absence of disease and it requires that the same check(s) be grown for many years consecutively.

If the breeding program data do not meet the requirements for neither the EBV nor the control population method, then the next best option would be to conduct an ERA trial. The advantage of conducting an ERA trial for estimation of realized *ΔG*_*t*_ is that the design of the experiment can be adjusted to achieve desired levels of error, precision, efficiency and correlation. For example, the number of lines used to represent a single year and the number of environments and replicates within an environment can be increased. The main disadvantage of this method is that it requires additional resources for designing and conducting the trial, and it relies on the availability of germplasm representative of the germplasm generated in each year of the breeding program. Most ERA trials that have been reported in the literature utilize very few varieties, less than one per year, to represent breeding germplasm developed over many years, possibly due to lack of seed sources of additional germplasm. In these cases, ERA trials are likely to be inadequate for estimating realized *ΔGt*. As an example, in this study, the ERA trial method with one line representing each year was just as accurate and less efficient as the simple phenotype method for estimating realized *ΔG*_*t*_ in the past 15 year time period. Thus the purpose of the ERA trial, which is to estimate realized *ΔG*_*t*_ more accurately compared to regression of average phenotype over years, was defeated entirely. An ERA trial with more lines sampled each year would have been more effective in this case.

### Indicators of realized ΔG_t_

Evaluating indicators of realized *ΔG*_*t*_ can be a useful first step prior to embarking on estimation of realized *ΔG*_*t*_ as it may indicate that realized *ΔG*_*t*_ > 0 is not expected, potentially eliminating unnecessary analyses or experiments attempting to estimate realized *ΔG*_*t*_. As demonstrated in this study, if accurate and complete pedigree records have been kept from the beginning of the breeding program, annual mean EqCg is associated with annual mean true breeding values. Thus, if advancement in mean EqCg is not observed over years, genetic gain would not be expected and attempting to estimate realized *ΔG*_*t*_ would be futile. Likewise, if average R/L ≯ 0, then realized *ΔG*_*t*_ also cannot be greater than zero. This study demonstrated that average R/L does in fact accurately predict the rate of realized *ΔG*_*t*_. The advantage of utilizing average R/L as an indicator of future realized *ΔG*_*t*_ is that it would enable routine monitoring of the selection decisions of breeding programs on a yearly basis so that potential problems or mistakes can be detected and corrected immediately rather than many years later when faulty selection decisions are finally reflected in estimates of realized *ΔG*_*t*_.

### Comparing breeding programs based on estimates of realized ΔG_t_

One of the most interesting findings of this study was that positive linear relationships between the true and estimated realized *ΔG*_*t*_ across breeding scenarios were not always observed. In fact, only the estimates of realized *ΔG*_*t*_ based on average R/L, control population, and ERA trial methods were at least moderately associated with true realized *ΔG*_*t*_ in the 15 year and 28 year time periods. Among these methods, the average R/L and the control population methods performed best.

The lack of association between estimates of realized *ΔG*_*t*_ and true *ΔG*_*t*_ was due to systematic error, also known as bias. Bias, the difference between true and estimated *ΔG*_*t*_, was associated with the testing scheme, heritability, and non-genetic trend. This makes it difficult to compare breeding programs effectively based on estimated realized *ΔG*_*t*_ because each breeding program will be subject to bias in a different way. For instance breeding program X may have a testing scheme that is associated with overestimation of realized *ΔG*_*t*_ while the testing scheme of breeding program Y is associated with underestimation of realized *ΔG*_*t*_. Even if these two breeding programs have the same true rate of realized *ΔG*_*t*_, breeding program X will appear to be performing better than breeding program Y. Even comparison of estimated realized *ΔG*_*t*_ between two different time periods within the same breeding program will be problematic if the breeding strategy differs between the two time periods.

While I do not recommend using the realized *ΔG*_*t*_ estimation or prediction methods presented in this paper for comparing breeding programs, itf would be relatively safe to use average R/L, control population, and ERA trial methods to make very general comparisons among breeding programs if the observed differences in estimated realized *ΔG*_*t*_ are very large. If breeding programs are compared based on estimated realized *ΔG*_*t*_, it will also be important to express estimated realized *ΔG*_*t*_ in units of genetic standard deviations because each breeding population will have a unique level of genetic variance that will have a major influence on the rate of realized *ΔG*_*t*_ when expressed in units of the trait of interest. Expressing *ΔG*_*t*_ in units of genetic standard deviations removes this effect.

If comparing breeding programs or breeding schemes is a major objective, methods other than those presented in this study should be utilized. The method proposed by Eberhart (1964), which has been used frequently in maize to evaluate realized *ΔG*_*t*_ from selection experiments, would be an effective method for this purpose. With this method, a sample of germplasm is taken each year or cycle, archived, and then tested years later in a common set of environments. While this is similar to the ERA trial method, it is better in that the sample of germplasm taken each year is relatively large and taken at random so that it can be considered a representative sample of the breeding population in a given year which should eliminate any association between sampling error and the breeding scheme. Another option would be to simply utilize simulation experiments to compare breeding schemes or programs. The major advantage of this approach are that the virtual breeding programs can be replicated and completed quickly at a low cost. Replication eliminates random noise that is inevitable due to random genetic drift, and the time and cost saved means that more breeding strategies can be evaluated and improved strategies can be implemented right away, avoiding a potentially major opportunity costs of not implementing an improved strategy once it is identified.

## Conclusion

Overall, this study showed that while its relatively easy to obtain estimates of realized *ΔG*_*t*_ using data generated from breeding programs or from ERA experiments, these estimates are often very inaccurate. Furthermore, the realized *ΔG*_*t*_ estimation methods evaluated in this study should not be used to compare breeding programs or to evaluate how changes made to breeding programs have affected realized *ΔG*_*t*_ because error in the estimates of realized *ΔG*_*t*_ is associated with the breeding scheme, non-genetic trend, and heritability. If the goal is simply to determine if a positive upward trend exists or to obtain a rough estimate of realized *ΔG*_*t*_, the control population, EBV, and ERA trial methods are recommended. The objectives, resources, and structure of the existing breeding program data that are available should be taken into account when selecting which of the three recommended methods, if any, should be utilized. No single method or or analysis pipeline be will appropriate for all situations, and for some breeding programs it may simply not be possible to obtain a meaningful estimate of realized *ΔG*_*t*_ with any method due to lack of appropriate data and seed stocks and/or because the breeding program was initiated only recently. In these cases, indicators of realized *ΔG*_*t*_ like expected *ΔG*_*t*_ and average EqCg would be useful to determine if the selection decisions made by the breeding program are consistent with a strategy that will lead to genetic gain over time.

## Acknowledgments

I would like to acknowledge Dr. Jean-Luc Jannink for giving constructive feedback on this research. I would also like to acknowledge Dr. George Kotch and Dr. Gary Atlin for our discussions about genetic gain estimation which exposed the need for more research on this topic. This work was funded by the Excellence in Breeding Platform (EiB) of the CGIAR, The CGIAR research program on Rice (Rice CRP), and the Transforming Rice Breeding (TRB) project funded by the Bill and Melinda Gates Foundation.

